# Dysconnectivity between auditory-cognitive network associated with auditory GABA and glutamate levels in presbycusis patients

**DOI:** 10.1101/2021.10.31.466279

**Authors:** Ning Li, Wen Ma, Fuxin Ren, Xiao Li, Fuyan Li, Wei Zong, Lili Wu, Zongrui Dai, Steve C.N. Hui, Richard A.E. Edden, Muwei Li, Fei Gao

## Abstract

Accumulating studies suggest an interaction between presbycusis (PC) and cognitive impairment, which may be explained by the cognitive-ear link to a large extent. However, the neurophysiological mechanisms underlying this link are largely unknown. Here, 51 PC patients and 51 well-matched healthy controls were recruited. We combined resting-state functional MRI and edited magnetic resonance spectroscopy to investigate changes of intra- and inter-network functional connectivity and their relationships with auditory gamma-aminobutyric acid (GABA) and glutamate (Glu) levels and cognitive impairment in PC. Our study confirmed the plastic model of cognitive-ear link at the level of the large-scale brain network, including the dysconnectivity within high-order cognitive networks and between the auditory-cognitive network and overactivation between cognitive networks dependent on hearing loss, which was closely related to the cognitive impairment of PC patients. Moreover, GABA and Glu levels in the central auditory processing were abnormal in patients with PC. Importantly, reduction of GABA-mediated inhibition plays a crucial role in a dysconnectivity between the auditory-cognitive network, which may be neurochemical underpinnings of functional remodeling of cognitive-ear link in PC. Modulation of GABA neurotransmission may enable the development of new therapeutic strategies for the cognitive impairment of PC patients.

## Introduction

Presbycusis (PC), defined as binaural symmetrical, progressive neurological deafness, is a condition of hearing impairment as aging. It is typically embodied as high-frequency hearing loss (HL), degraded speech comprehension and perception especially in noisy environments ^1^. Obviously, hearing loss is the manifestation of human aging in auditory organs, which is usually related to the physiological decline of inner ear cells and auditory cortex. However, the decline of speech understanding in PC patients, especially in noisy environment, seems far beyond the function range of auditory cortex. Besides, accumulating evidence proved that PC was related to accelerate cognitive decline, incident cognitive impairment (CI) and dementia ^2, 3,4^. Thus, it was convincing that in addition to the impairment of inner ear function, PC also shows a dysfunction of the central auditory pathways, called as the central auditory processing disorder (CAPD). Some studies suggested that compared to peripheral HL, CAPD is more likely to cause cognitive decline ^5^. Based on these studies, the cognitive-ear link was then proposed, which suggested that in addition to the ear and auditory cortex, there were other cognition-related cortices also participating in the auditory information processing ^6, 7^. Furthermore, PC may cause change of the original cognitive-ear link and produce extensive structural and functional remodeling. However, their exact neuropathogical basis are largely unknown.

Previous magnetic resonance imaging (MRI) research have demonstrated that the auditory cortex and cognitive related cortex participate in CI of patients with PC ^8–10^. As reported, PC was associated with the gray matter volume decrease in primary auditory cortex (i.e., Heschl’s gyrus, HG) and other temporal lobe structures ^11–13^. Another structural MRI study using surface-based morphometry confirmed that PC patients presented with gray matter (GM) atrophy in several auditory cortical areas and nodes of the default mode network (DMN), such as the bilateral precuneus and right posterior cingulate cortex (PCC) ^14^. Moreover, a negative correlation was found between the scores on the Trail-Making Test (TMT), which evaluates executive control, and the cortical volume of the right PCC in PC patients ^14^. Abnormal amplitude of low-frequency fluctuation (ALFF) in the HG, dorsolateral prefrontal cortex (dlPFC), frontal eye field and key nodes of the DMN, were also observed in PC patients, which associated with specific cognitive performance, such as verbal learning and memory ^15^. These findings suggest that the occurrence of PC seems to initiate a series of related cascade reactions, which might lead to cross-modal plasticity rather than the structural and functional changes in a single cerebral cortex.

By studying the low frequency fluctuations of BOLD signals in different brain regions at resting state, function MRI (fMRI) can reflect the neurophysiological relationship between spatially distant brain regions and evaluate the functional synergy between brain regions ^16^. Recent resting-state functional connectivity (rsFC) studies have confirmed stronger FC between the dlPFC and the poster dorsal stream of auditory processing, weaker FC between the PCC and key nodes of the DMN, as well as weaker FC between the hippocampus and insula, which were related with specific CI in PC patients ^15^. Although altered rsFC between two cerebral cortices were revealed by this approach, the changes of intra- and inter-network connectivity in the brain cannot be explained. Thus, a better understanding of the neuropathogical mechanism of the cognitive-ear link in PC would require investigations at the level of the brain network. Independent component analysis (ICA) is a data-driven method that can divide MRI data into some spatially independent components without a priori hypothesis. After separate the interference components such as heartbeat, respiration, head movement, etc., the remaining spatially specific components can form a network, usually called resting-state networks (RSNs). It can assessment the synchronization and the interactions among multiple networks, thus overcomes the shortcomings of time-dependent analysis ^17^. Its reliability and validity have been verified in the study of a series of cognitive disorders ^18–21^.

As the main excitatory and inhibitory neurotransmitters, glutamate (Glu) and gamma-aminobutyric acid (GABA) play an important role in maintaining the balance of central auditory processing ^22^. Moreover, GABA is also participated in neural plasticity and synchronous network oscillation, which is very important for effective information processing and normal cognitive function ^22–24^. A magnetic resonance spectroscopy (MRS) study indicated decreased auditory Glu levels in PC patients when compared with the young healthy control group. However, GABA levels are difficult to measure using conventional MRS due to the relatively low level and the resonance of GABA usually overlaps with other metabolites ^25^. A recent edited MRS called MEGA-PRESS which can separate GABA signals from other metabolisms realizes the quantitative measurement of GABA levels ^26–28^. Then, we reported decreased auditory GABA levels in PC patients using MEGA-PRESS method ^8^. In addition, another MEGA-PRESS study found that decreased GABA levels were associated with cognitive decline in the elderly, and speculated that GABA levels may play an important role in neural dedifferentiation ^29^. Meanwhile, the generation of BOLD fMRI signal depends on the excitatory glutamatergic projection neurons, while GABAergic interneurons may indirectly regulate the BOLD signal through inhibitory feedback signal in microcircuit. Importantly, there are key regulation functions for glutamatergic and GABAergic systems in functionally connected cortical circuits. Stagg and his colleagues demonstrated that primary motor GABA levels were associated with FC intensity in the motor network in healthy participants ^30^. Considering altered GABA and Glu levels in PC patients as measured using MRS, these neurotransmitter systems are likely to be related with the alterations of functional network connectivity (if it is indeed changed).

In this study, therefore, we applied MEGA-PRESS with macromolecule suppression and PRESS to examine the change in auditory GABA and Glu levels in PC patients. Then, the changes of FC at the level of brain network were investigated using an ICA-based RSNs analysis. Based on previous findings, the interactions among FC strengths, levels of metabolites and cognitive function were evaluated to test our hypothesis: (1) whether intra- and inter-network FC strengths were altered in PC patients; (2) whether auditory GABA or Glu levels was correlated with FC strengths in PC patients; (3) whether cognitive function was associated with auditory GABA or Glu levels in PC patients; (4) whether cognitive function was correlated with FC strengths in PC patients. This study might enhance our understanding on the plasticity of cognitive-ear link at the level of RSNs and neurochemical underpinnings of this plasticity in PC patients.

## Material and methods

### Subjects

Fifty-one PC patients (28 males/23 females, mean age, 65.16 ± 2.43 years) and the same number of normal hearing (NH) controls (21 males/30 females, mean age, 64.67 ± 1.67 years) matched with age, gender and education were enrolled in this research (Table 1). All subjects were right-handed and native Mandarin Chinese speakers with age ≥ 60 years. We used pure tone average (PTA) to evaluate HL at the thresholds of 0.5, 1, 2, and 4 kHz, and PTA = 25 dB HL was taken as the critical value ^31^. Specifically, for PC group, the inclusion criteria were PTA > 25 dB HL in the better hearing ear while it was PTA ≤ 25 dB HL for the NH group. Details of the exclusion criteria are provided in the supplemental material. This research was approved by the Shandong University Institutional Review Board, and the written informed consent was obtained from each subject.

**Table 1.** Demographic and clinical data of the NH group and PC group.

| **Characteristics** | **NH Group** | **PC Group** | ***p*-value** |
| --- | --- | --- | --- |
|  | (n = 51) | (n = 51) |  |
| Age (years) | 64.67 ± 1.67 | 65.16 ± 2.43 | 0.238 |
| Gender (male/female) | 21/30 | 28/23 | 0.165 |
| Disease duration (years) | -- | 5.80 ± 4.90 | -- |
| Education (years) | 11.43 ± 2.56 | 10.31 ± 4.37 | 0.119 |
| Anxiety | 3.61 ± 3.38 | 3.04 ± 3.19 | 0.384 |
| Depression | 3.33 ± 3.52 | 3.84 ± 3.89 | 0.489 |
| Alcohol abuse (yes/no) | 2/49 | 4/47 | 0.674 |
| Smoking (yes/no) | 4/47 | 9/42 | 0.138 |
| Hyperlipemia (yes/no) | 9/42 | 9/42 | 1.000 |
| Hypertension (yes/no) | 20/31 | 27/24 | 0.164 |
| Diabetes (yes/no) | 8/43 | 8/43 | 1.000 |

**Notes:** The data are presented as means ± standard deviations. levels of anxiety and depression were assessed according to the Hospital Anxiety and Depression Scale (HADS).
**Abbreviations:** NH, normal hearing; PC, presbycusis.

### Auditory Assessment

The pure tone threshold was assessed via a clinical audiometer (Madsen Electronics Midimate 622) coupled with TDH-39P telephonic headphones for each ear separately at frequencies of 0.125, 0.25, 0.5, 1, 2, 4, and 8 kHz. Speech detection was assessed using speech reception threshold (SRT), following the guidelines recommended by the American Speech-language Hearing Association. Details of the auditory assessment are provided in the supplemental material.

### Neuropsychological Assessment

Neuropsychological tests were performed for each subject in a certain order within about 1 hour. First, Montreal Cognitive Assessment (MoCA) ^32, 33^ was performed to test subject’s general cognitive function. Then, Auditory Verbal Learning Test (AVLT, Chinese version) ^34^, Stroop Color-Word Interference Test (Stroop) ^35^, Symbol Digit Modalities Test (SDMT) ^36^, TMT-A and TMT-B ^37^ were performed to assess auditory learning and memory, attention, information processing speed, and executive function, respectively. Finally, Hospital Anxiety and Depression Scale were applied to assess subject’s emotion ^38^.

### MRI acquisition

All subjects were examined under a 3.0-Tesla MR (Philips Achieva TX, Best, Netherlands) with an 8-channel head coil. The functional data were obtained using a gradient EPI sequence (TR/TE = 2000/35 ms, FOV = 240 × 240 mm^2^, in-plane resolution = 3.75 × 3.75 mm^2^, 35 slices with a 4-mm slice thickness, and 240 dynamics.

The structural data were obtained using a 3D T1-weighed sequence (TR/TE = 8.1/3.7 ms, FOV = 240 × 240 mm^2^, 160 slices with a 1-mm slice thickness, and 1 mm isotropic voxels). We selected bilateral auditory regions centered on HG as the volumes of interest (VOIs) with a 40 × 30 × 20-mm^3^ size, following the criteria recommended by Rojas et al. ^39^ (as shown in Fig. 1). The GABA-edited data were obtained using a Macromolecules (MM)-suppressed MEGA-PRESS sequence ^40^ (TR/TE=2000/80 ms, bandwidth = 2000 Hz, ‘‘ON/OFF’’ editing pulses = 1.9/1.5 ppm, 320 averages), while the symmetrical-suppression method was used in suppressing MM signal ^40^. According to differences between the ON and OFF spectra, we can obtain the results of edited-GABA. The Glu data were obtained using a PRESS sequence at the same VOI (TR/TE=2000/35 ms, bandwidth = 2000 Hz, 64 averages). The unsuppressed water data were obtained using a shorter measurement with 8 averages.

**Figure 1.**
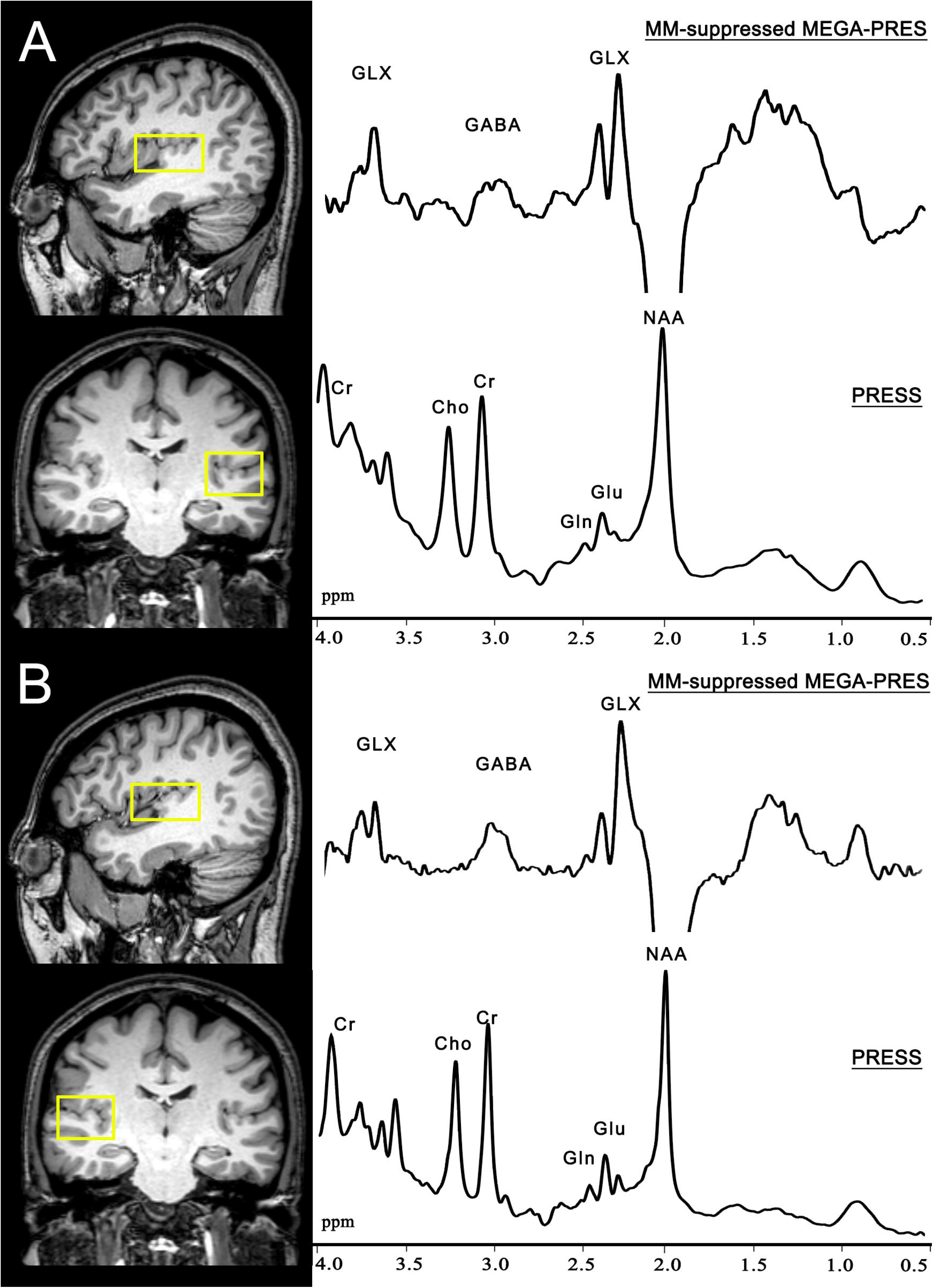
Representative Macromolecules (MM)-suppressed MEGA-PRESS spectra are shown from the volumes of interests (4 × 3 × 2 cm^3^) centered on the Heschl’s gyrus in the left auditory region (A) and the right auditory region (B), respectively.

### Functional Data Preprocessing

The functional data were preprocessed by the Data Processing & Analysis for Brain Imaging (DPABI) V5.1 toolbox ^41^. The processing steps were as follows: (1) excluded the first 10 volumes out of all 240 dynamics; (2) corrected for the slice-timing and realigned for head motion correction; (3) co-registered T1-weighted and functional data, and segmented them into GM, white matter (WM) and cerebrospinal fluid (CSF); (4) spatially normalized the co-registered functional data to the standard space; (5) applied nuisance covariate regression with the Friston 24 head motion parameters ^42^ and CSF signal, and then smoothed the images with an isotropic Gaussian kernel (FWHM = 4 mm); (6) performed a linear detrending and temporal filtering (0.01-0.08 Hz).

### Independent Component Analysis

For independent components (ICs), the Group ICA of fMRI Toolbox (GIFT) software (http://icatb.sourceforge.net/) was applied as follows: (1) data reduction at the individual level was performed by using a two-step principal component analysis (PCA), and 40 ICs were extracted for each subject; (2) extracting ICs by an Infomax ICA algorithm and generating a final set of ICs; (3) reconstructing the time courses and spatial maps according to the composition of each subject data set and the results of data dimensionality reduction. (4) All results were converted into z-scores to display, and that best matched the RSNs template were selected for further evaluation.

### Functional Network Connectivity analysis

Finally, 12 out of 40 ICs were chosen as the focus for later FC analysis. They are spatially independent but exhibit significant dependencies in temporal aspect. The FC graph was extracted and transformed into z-value graph by Fisher’s method before analysis. The differences of intra-network FC between groups were evaluated by a two-sample t-test, taken the age, gender and education as covariates. The correction for multiple comparisons was applied using an FDR approach with *p* < 0.05 and cluster size > 5 voxels. Comparisons between-group of all 12 × 12 inter-network FC matrices were also conducted by a two-sample t-test, taken the age, gender and education as covariates. The correction for multiple comparisons was applied using an FDR approach with *p* < 0.05.

### MRS data analysis

The MM-suppressed MEGA-PRESS and PRESS data were processed using Gannet 3.1 ^43^ and LCModel (version 6.3-1 M) ^44^. Gannet applied 3 Hz line broadening and Gaussian curve to fit GABA peaks at 3 ppm. Then, Gannet co-registered VOIs to the 3D T1-weighted data and segmented VOIs into GM, WM and CSF. Spectra with fitting errors of GABA < 20% and Cramer-Rao lower bounds (CRLB) of Glu < 20% were chosen for further investigation.

### Statistical analysis

Comparisons between-group of age, education, anxiety/depression scores, MRS data, and cognitive function scores were conducted by a two-sample t-test. Other clinical data, such as gender and hyperlipemia, were evaluated by a two-sample Chi-square test.

Taken the age, gender and education as covariates, we applied partial correlation analysis to explore the interactions among FC strengths, levels of GABA and Glu, cognitive function in PC and NH groups.

A random forest method was used to evaluate the importance of the imaging indexes in predicting the clinical features. Here we used two widely accepted criteria ‘IncMSE%’ and ‘IncNodePurity’ for feature assessment. IncMSE% equals the increased mean of the squared error while the feature is rearranged randomly, and IncNodePurity means the percentage of increased Gini parameter after the random rearrangement. To fit the tendency of MOCA, AVLT, SDMT, Stroop, TMT-A, and TMT-B, 76 features were used to make the prediction. The random forest model was built by R.3.6.0 with the random forest package based on default parameters.

## Results

### Comparisons of demographic and clinical data

No significant differences were found between-group in age, gender, education, anxiety/depression scores, AVLT score, and hyperlipemia, etc. (all *p* > 0.05, Table 1 and 2). Compared with the NH group, PC patients had poor performance in cognitive function, such as MoCA, SDMT, Stroop, TMT-A, and TMT-B tests (all *p* < 0.01, Table 2), and poor performance in auditory function, such as PTA and SRT (all *p* < 0.001, Table 2). All subjects had normal middle ear function with a type-A tympanometry curve. The average pure tone threshold of bilateral ears at frequencies of 0.125-8 kHz was shown in Fig. 2

**Table 2.**
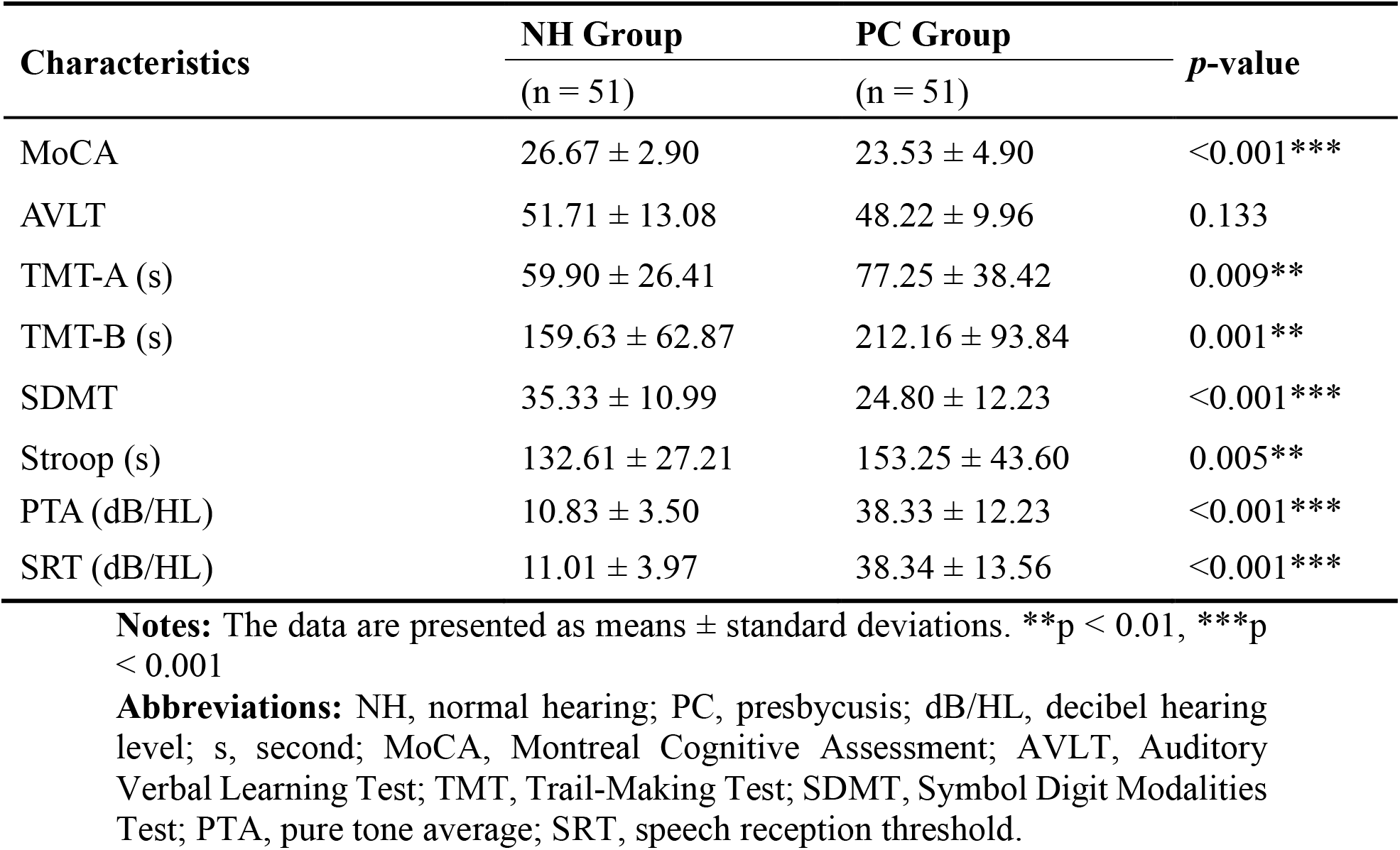
Auditory and cognitive function data of the NH group and PC group.

| **Characteristics** | **NH Group** | **PC Group** | ***p*-value** |
| --- | --- | --- | --- |
|  | (n = 51) | (n = 51) |  |
| MoCA | 26.67 ± 2.90 | 23.53 ± 4.90 | <0.001*** |
| AVLT | 51.71 ± 13.08 | 48.22 ± 9.96 | 0.133 |
| TMT-A (s) | 59.90 ± 26.41 | 77.25 ± 38.42 | 0.009** |
| TMT-B (s) | 159.63 ± 62.87 | 212.16 ± 93.84 | 0.001** |
| SDMT | 35.33 ± 10.99 | 24.80 ± 12.23 | <0.001*** |
| Stroop (s) | 132.61 ± 27.21 | 153.25 ± 43.60 | 0.005** |
| PTA (dB/HL) | 10.83 ± 3.50 | 38.33 ± 12.23 | <0.001*** |
| SRT (dB/HL) | 11.01 ± 3.97 | 38.34 ± 13.56 | <0.001*** |

**Notes:** The data are presented as means ± standard deviations. \*\* $p < 0.01$ , \*\*\* $p < 0.001$
**Abbreviations:** NH, normal hearing; PC, presbycusis; dB/HL, decibel hearing level; s, second; MoCA, Montreal Cognitive Assessment; AVLT, Auditory Verbal Learning Test; TMT, Trail-Making Test; SDMT, Symbol Digit Modalities Test; PTA, pure tone average; SRT, speech reception threshold.

**Figure 2.**
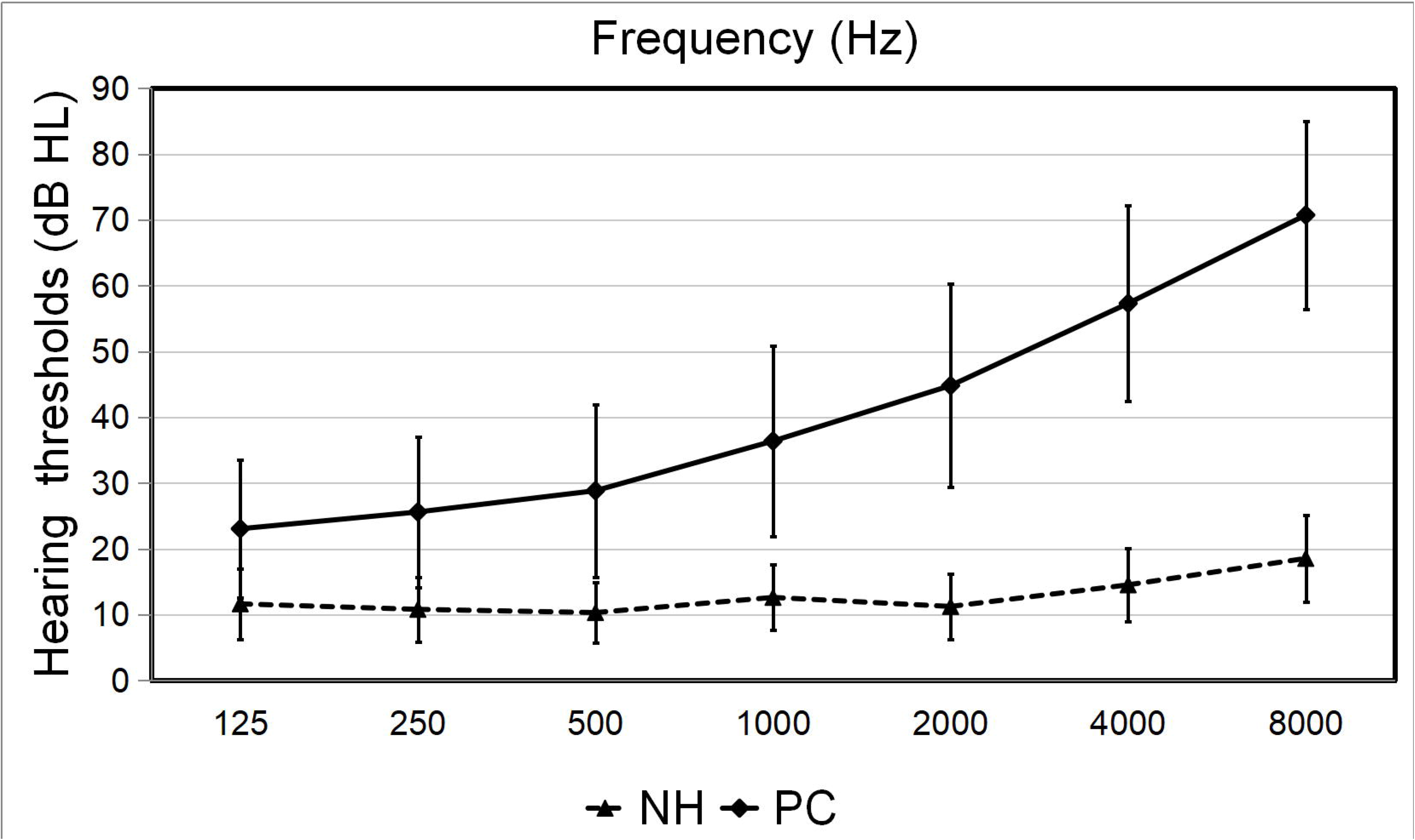
The average pure tone threshold of bilateral ears at frequencies of 0.125-8 kHz in the PC and NH groups.

### Comparisons of MRS data in the auditory region

No significant differences were found in the GM or WM fractions between groups within the left auditory regions (AR) and right AR (all *p* > 0.05, Table 3). Some values of GABA and Glu measurement were excluded from analysis based on the fitting error and CRLB as described in the methods (GABA: left AR, 1 control and 2 patients; right AR: 2 controls. Glu: left AR, 2 patients). No significant difference was observed in the fitting error and CRLB between groups in the left AR and right AR (all *p* > 0.05, Table 3). Compared to the NH group, significantly lower GABA levels were found in the right AR in patients with PC (0.91 ± 0.19 iu vs 1.08 ± 0.31 iu, *p* = 0.002), whereas no significant differences were found in the left AR between PC and NH groups (0.92 ± 0.31 iu vs 1.22 ± 1.72 iu, *p* = 0.234). Besides, decreased Glu levels were found in bilateral ARs in patients with PC (left AR: 6.07 ± 0.88 iu vs. 6.53 ± 0.57 iu, *p* = 0.003, right AR: 6.54 ± 0.72 vs 6.85 ± 0.52 iu, *p* = 0.015, Table 3).

**Table 3.** MRS data of the NH group and PC group.

| Characteristics | NH Group | PC Group | *p*-value |
| --- | --- | --- | --- |
| **Left auditory region** |  |  |  |
| GABA levels (iu) | 1.22±1.72 | 0.92±0.31 | 0.234 |
| GABA fitting errors | 9.16±2.81 | 9.44±3.06 | 0.639 |
| GMF (%) | 52.24±5.06 | 51.99±3.71 | 0.781 |
| WMF (%) | 32.01±8.33 | 30.71±5.53 | 0.357 |
| Glu levels (iu) | 6.53±0.57 | 6.07±0.88 | 0.003** |
| Glu CRLB | 5.75±0.74 | 5.90±1.16 | 0.437 |
| **Right auditory region** |  |  |  |
| GABA levels (iu) | 1.08±0.31 | 0.91±0.19 | 0.002** |
| GABA fitting errors | 9.54±2.99 | 9.90±3.16 | 0.554 |
| GMF (%) | 53.29±4.87 | 52.82±3.80 | 0.588 |
| WMF (%) | 31.27±8.04 | 30.86±4.94 | 0.757 |
| Glu levels (iu) | 6.85±0.52 | 6.54±0.72 | 0.015* |
| Glu CRLB | 5.55±0.58 | 5.78±1.21 | 0.213 |

**Notes:** The data are presented as mean ± standard deviations. \* $p < 0.05$ , \*\* $p < 0.01$ .
**Abbreviations:** MRS, magnetic resonance spectroscopy; iu, institutional units; PC, presbycusis; NH, normal hearing; GABA, gamma-aminobutyric acid; Glu, glutamate; GMF, gray matter fractions; WMF, white matter fractions; CRLB, Cramer-Rao lower bound.

### Resting-state networks

12 ICs were acquired by using group ICA approach. These 12 RSNs are demonstrated in Fig. 3, including the anterior default mode network (aDMN; IC 1), posterior default mode network (pDMN; IC 20), default mode network (DMN; IC 38), visual network (VN; IC 13), auditory network (AN; IC 10), sensorimotor network (SMN; IC 33), left executive control network (lECN; IC 35), right executive control network (rECN; IC 36), basal ganglia network (BGN; IC 34), salience network I (SN I; IC 5), SN II (IC 37), and dorsal attention network (DAN; IC 28).

**Figure 3.**
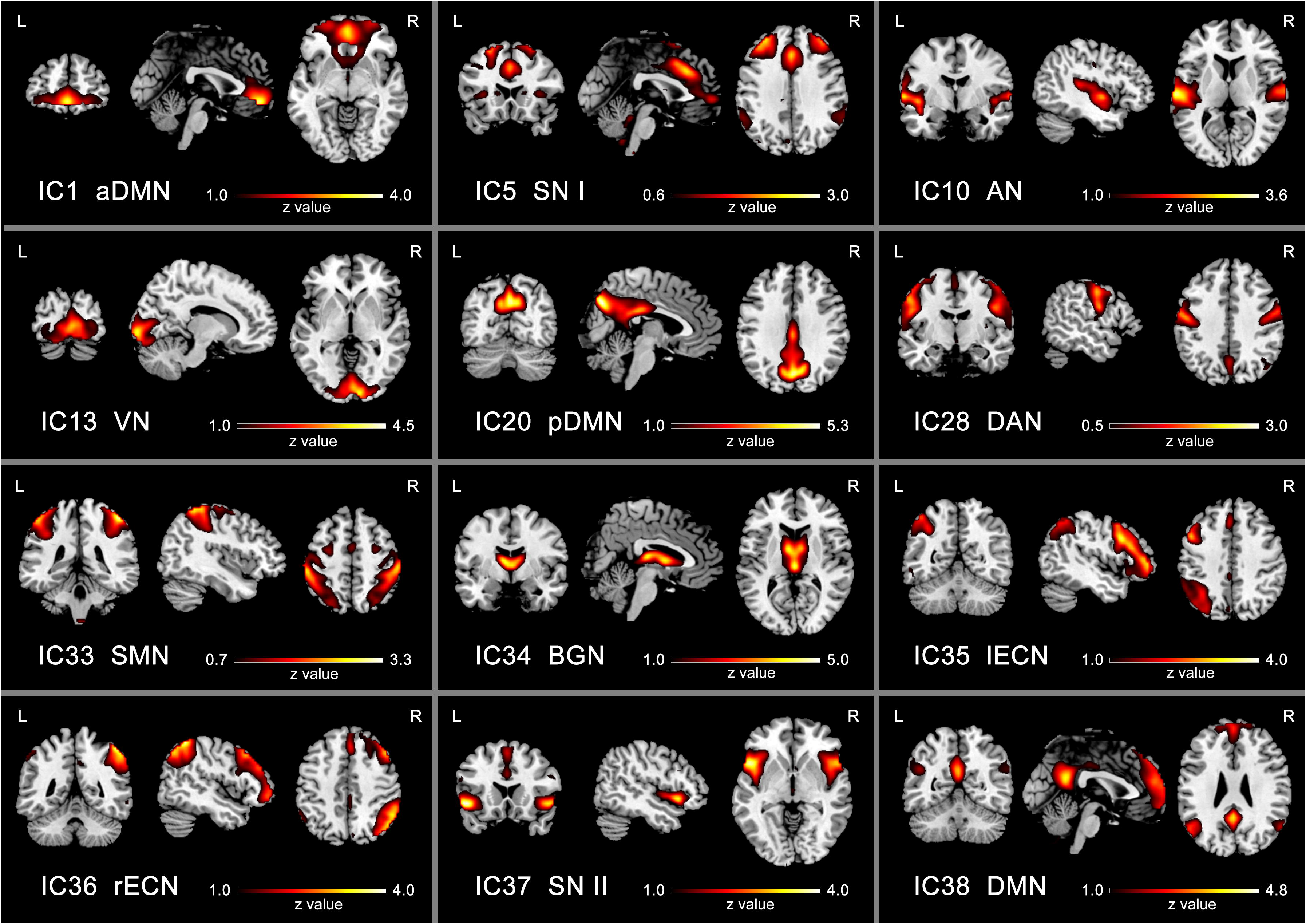
Spatial distribution of 12 intrinsic resting-state networks (RSNs) determined by independent component analysis (ICA). The colormaps represent the z-values.

### Comparisons of Intra-network Functional Connectivity

Intra-network FC changes were found in pDMN, DMN, lECN and rECN. Compared to the NH group, PC patients showed significantly decreased FCs in bilateral precuneus within pDMN and DMN, and decreased FCs in bilateral PCC within DMN. Compared to the NH group, PC patients showed significantly decreased FCs in left superior frontal cortex (SFC) and left inferior parietal lobule (IPL) within lECN, and decreased FCs in right dorsolateral prefrontal cortex (dlPFC) and right SFC within rECN (FDR corrected, *p* < 0.05, cluster size > 5 voxels), as seen in Table 4 and Fig. 4.

**Table 4.** Brain regions with significant differences in the intra-network functional connectivity between the PC group and NH group.

| RSNs | Brain region | Brodmann area | MNI coordinates |  |  | T value | Cluster size |
| --- | --- | --- | --- | --- | --- | --- | --- |
|  |  |  | x | y | z |  |  |
| pDMN | L posterior cingulate cortex | 23 | 3 | -21 | 24 | 6.2247 | 15 |
| pDMN | R posterior cingulate cortex | 23 | 3 | -21 | 24 | 6.2247 | 15 |
| pDMN | L precuneus | 7 | -15 | -69 | 36 | 5.4475 | 97 |
| pDMN | R precuneus | 7 | -15 | -69 | 36 | 5.4475 | 52 |
| DMN | L precuneus | 7 | -18 | -54 | 15 | 7.2241 | 14 |
| DMN | R precuneus | 7 | 21 | -51 | 15 | 7.7276 | 18 |
| IECN | L superior frontal cortex | 10 | 0 | 27 | 36 | 4.49 | 10 |
| IECN | L inferior parietal lobule | 39 | -36 | -57 | 42 | 4.968 | 20 |
| rECN | R superior frontal cortex | 10 | 21 | 24 | 54 | 5.03 | 11 |
| rECN | R dorsolateral prefrontal cortex | 9 | 39 | 27 | 36 | 4.3899 | 12 |

**Notes:** FDR corrected $p < 0.05$ , cluster size $> 5$ voxels.
**Abbreviations:** PC, presbycusis; NH, normal hearing; RSNs, resting-state networks; MNI, Montreal Neurological Institute; DMN, default mode network; pDMN, posterior default mode network; IECN, left executive control network; rECN, right executive control network; L, left; R, right.

**Figure 4.**
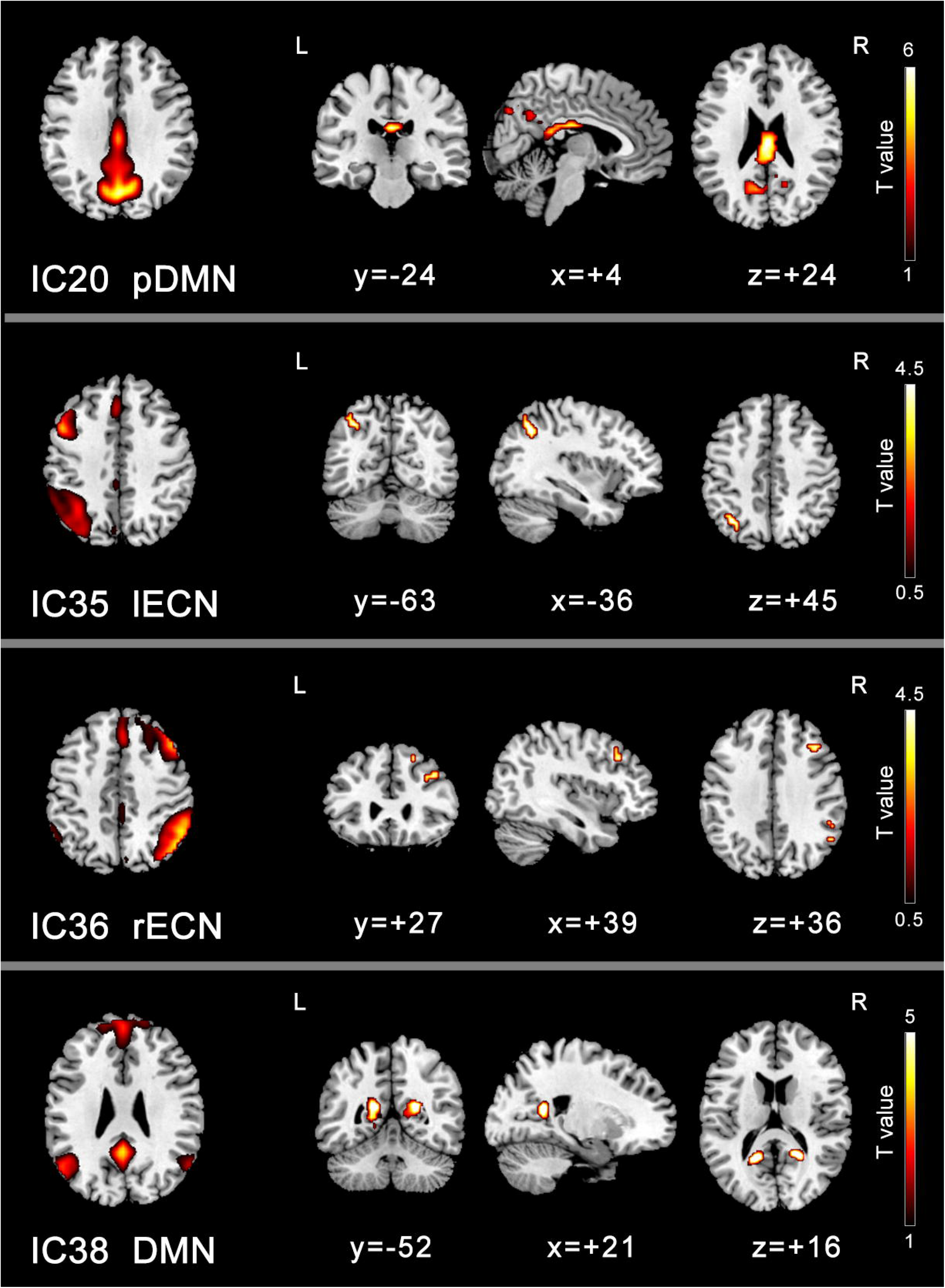
Group difference distributions of intra-network FC between PC group and NH group. Significantly decreased FC within posterior default mode network (pDMN), left executive control network (lECN), right executive control network (rECN) and default mode network (DMN) are found in the PC group (*p* < 0.05, FDR corrected).

### Comparisons of Inter-network Functional Connectivity

Compared to the NH group, PC patients showed obviously decreased inter-network FC between AN and DMN, aDMN and SN I, SN I and SN II, SN I and VN, SN I and BGN, whereas obviously increased FC between aDMN and SN II (FDR corrected, *p* < 0.05, Fig.5). Additionally, PC patients showed increased trend of inter-network FC between DMN and DAN, aDMN and pDMN (Fig.5).

**Figure 5.**
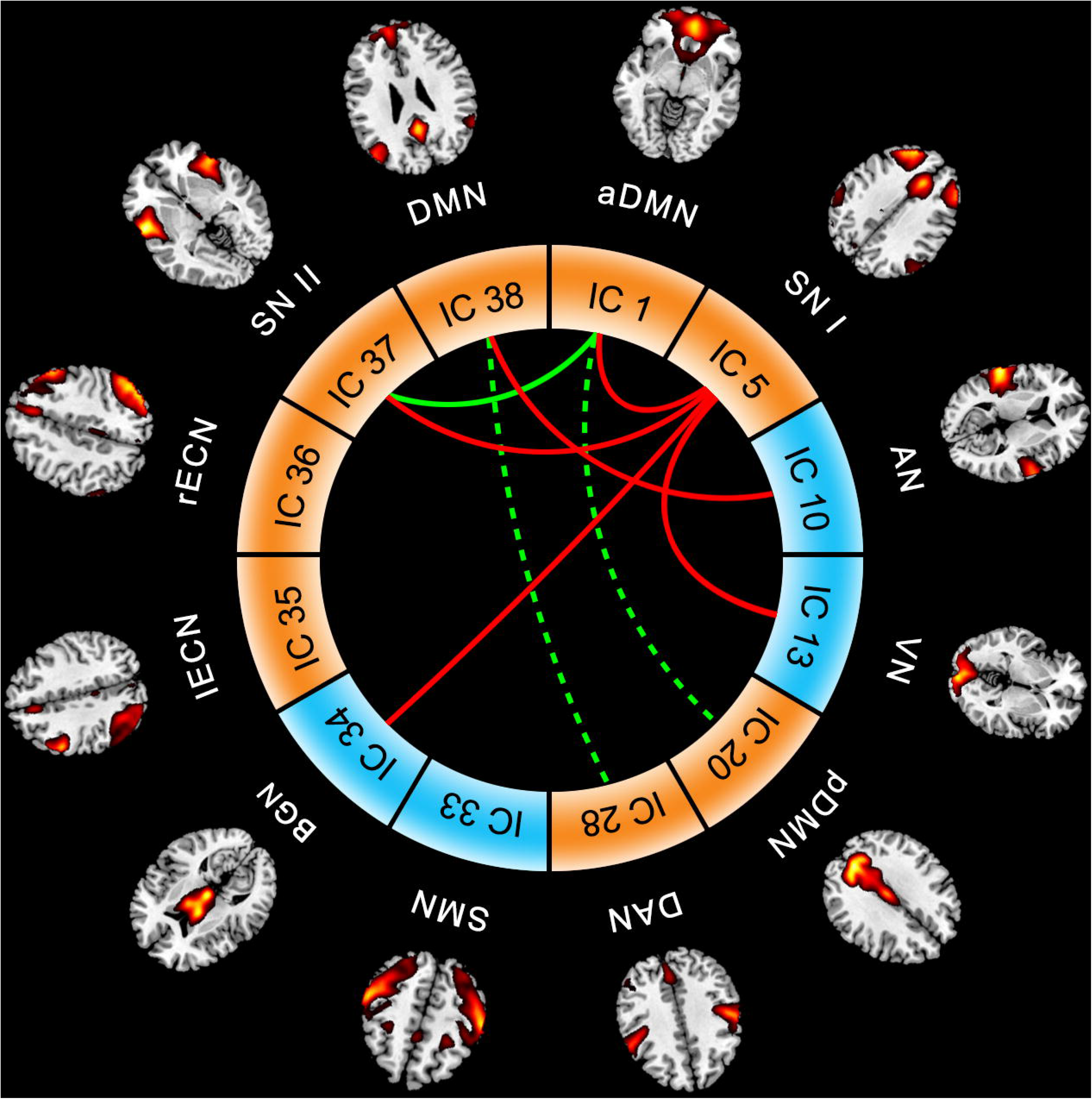
Group difference in the inter-network FC between the PC group and NH group. The lines connect the arc pairs represent significant differences in the FC between corresponding resting-state networks pairs (*p* < 0.05, corrected by FDR). Red lines denote significantly decreased inter-network FC; green lines denote significantly increased inter-network FC; green dotted lines denote increased trend of inter-network FC in the PC group.

### Relationships between FC strengths and clinical performance

For PC group, intra-network FCs of left SFC were positively correlated the MoCA (*r* = 0.288, *p* = 0.047) and SDMT (*r* = 0.484, *p* < 0.001), as seen in Fig. 6. The trend of correlations was found between FCs of right dlPFC and SDMT (*r* = 0.282, p = 0.052), and between FCs of left precuneus and PTA (*r* = −0.285, *p* = 0.050), as seen in Fig. 6. For PC group, inter-network FCs between aDMN and SN II were positively correlated the PTA (*r* = 0.349, *p* = 0.015), FCs between AN and DMN were negatively correlated the TMT-A (*r* = −0.389, *p* = 0.006), FCs between AN and pDMN were positively correlated the MoCA (*r* = 0.326, *p* = 0.024) and AVLT (*r* = 0.294, *p* = 0.043), as seen in Fig. 7. However, no relationships were found between FCs and clinical performance in the NH group.

**Figure 6.**
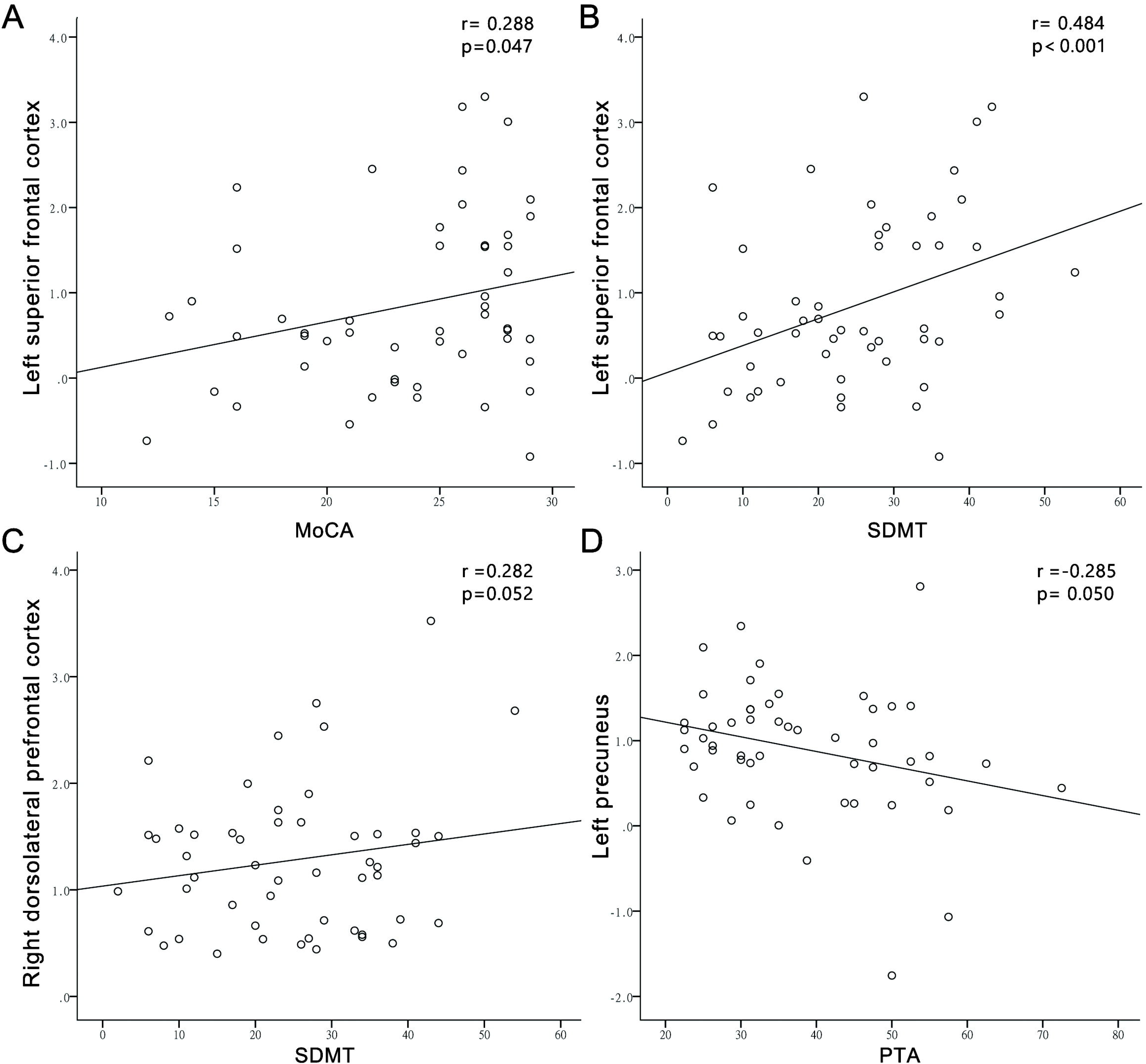
Relationships between intra-network FC strengths (FCs) and clinical performance in the PC group. The left SFC were positively correlated the MoCA **(A)** and SDMT **(B)**. The trend of correlations was found between FCs of right dorsolateral prefrontal cortex and SDMT (**C**), and between FCs of left precuneus and PTA (D).

**Figure 7.**
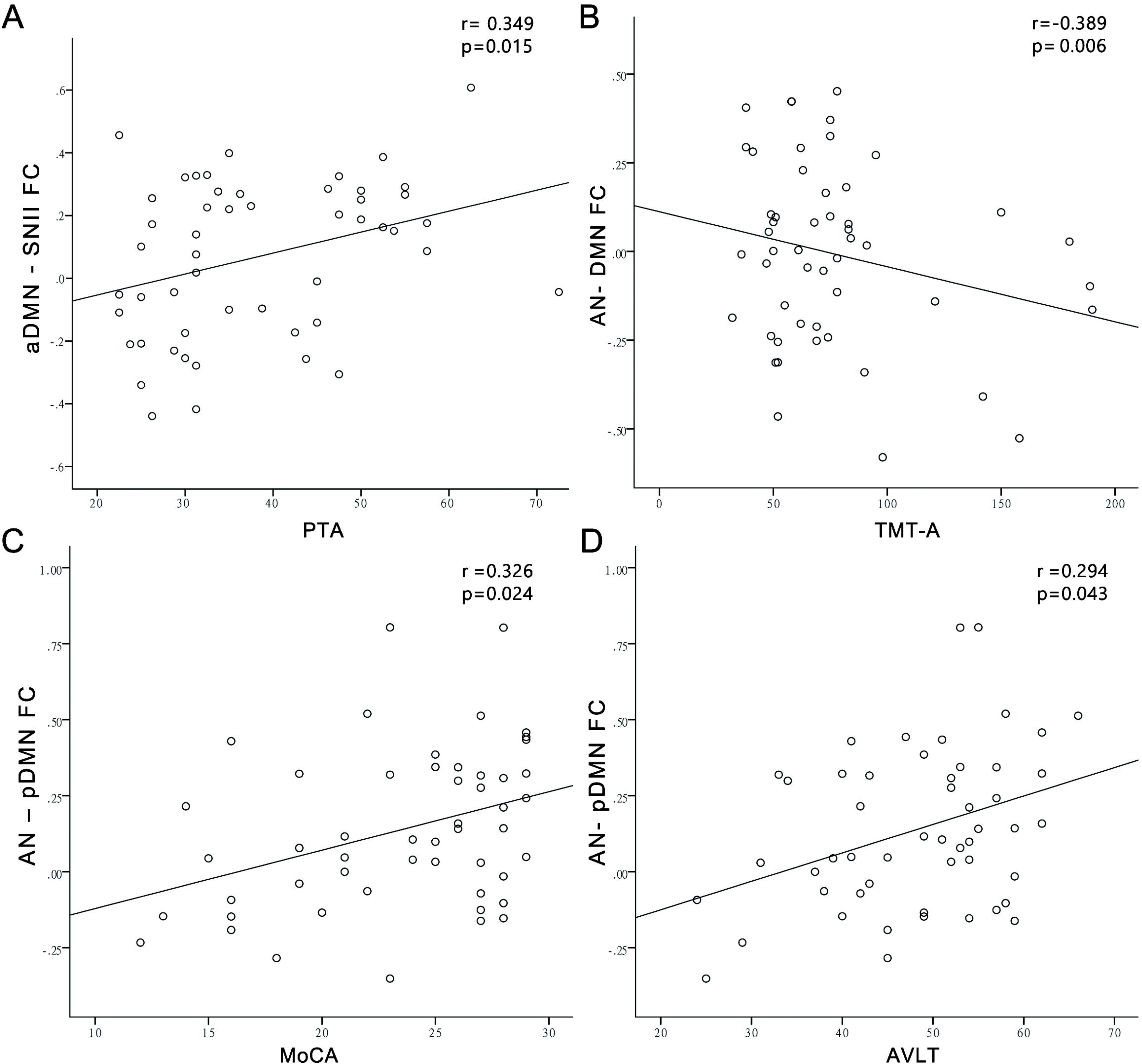
Relationships between inter-network FC strengths (FCs) and clinical performance in the PC group. FCs between aDMN and SN II were positively correlated the PTA (**A**), FCs between AN and DMN were negatively correlated the TMT-A (**B**), FCs between AN and pDMN were positively correlated the MoCA (**C**) and AVLT (**D**). aDMN, anterior default mode network; pDMN, posterior default mode network; DMN, default mode network; AN, auditory network.

### Relationships of the GABA/Glu levels and FC strengths

For PC group, GABA levels in the right AR were positively correlated with inter-network FCs between AN and DMN (*r* = 0.302, *p* = 0 .037) and AN and aDMN (*r* = 0.358, *p* = 0.013), as seen in Fig. 8. Glu levels in the right AR were positively correlated with FCs between AN and rECN (*r* = 0.333, *p* = 0.021), AN and SN II (*r* = 0.484, *p* = 0.000) and AN and pDMN (*r* = 0.374, *p* = 0 .009), while Glu levels in the left AR were positively correlated with FCs between AN and rECN (*r* = 0.337, *p* = 0.022), as seen in Fig. 8. However, no relationships were found between GABA or Glu levels and FCs in the NH group.

**Figure 8.**
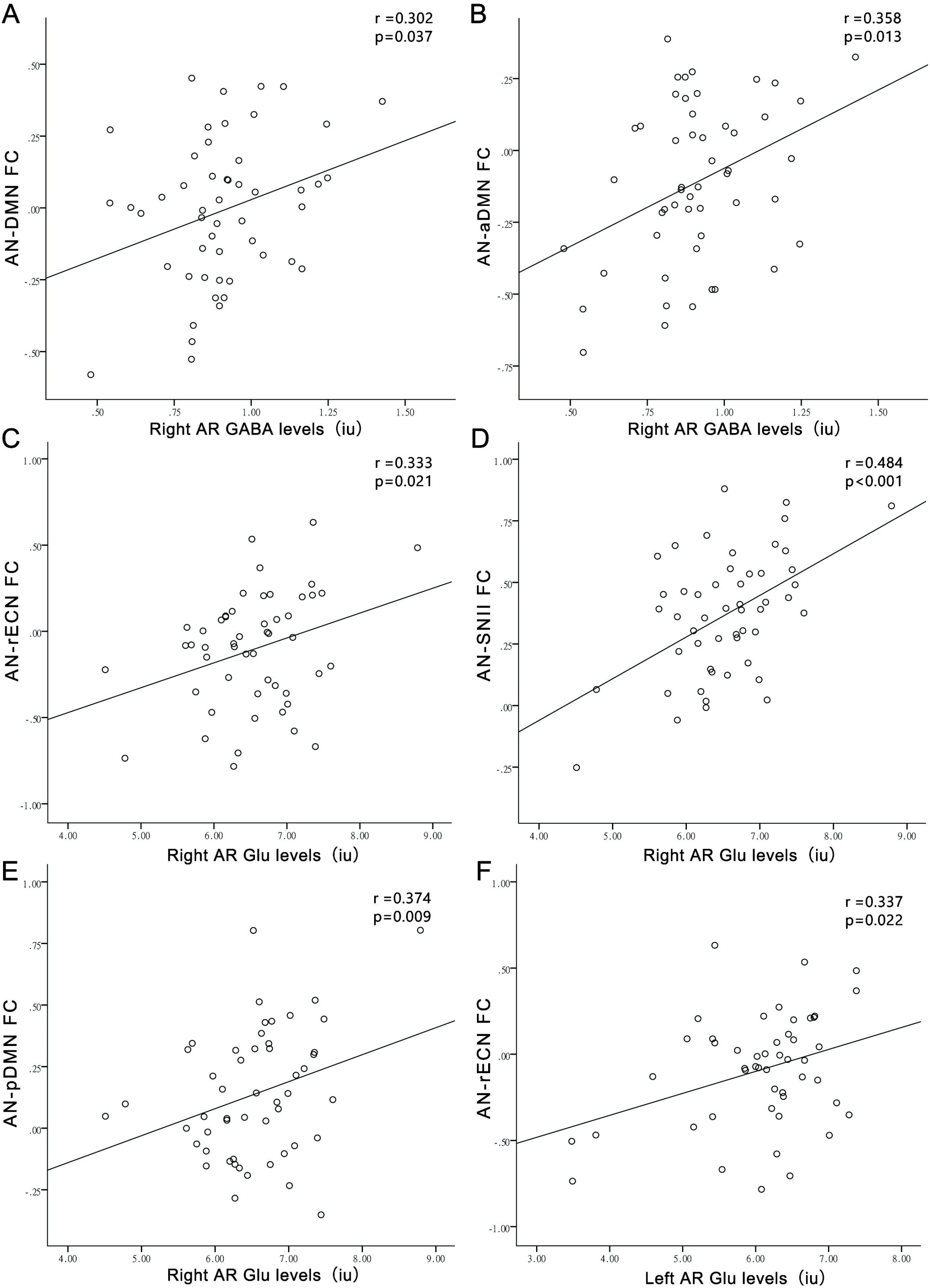
Relationships of the auditory region (AR) GABA/Glu levels and inter-network FC strengths (FCs) in the PC group. The right AR GABA levels were positively correlated with FCs between AN-DMN **(A)**, and between AN-aDMN **(B)**. The right auditory Glu levels correlated positively with FCs between AN-rECN **(C)**, and AN-SN II **(D)**, and AN-pDMN **(E)**. The left AR Glu levels correlated positively with FCs between AN-rECN **(F)**. aDMN, anterior default mode network; pDMN, posterior default mode network; DMN, default mode network; AN, auditory network; lECN, left executive control network; rECN, right executive control network; SN, salience network.

### Relationships between GABA/Glu levels and clinical performance

For PC group, GABA levels in the right AR were positively correlated with MoCA (*r* = 0.331, *p* = 0.022), while the trend of correlation was found between GABA levels in the right AR and AVLT (*r* = 0.246, *p* = 0.092), as seen in Fig. 9. However, no relationships were found between GABA/Glu levels and clinical performance in the NH group.

**Figure 9.**
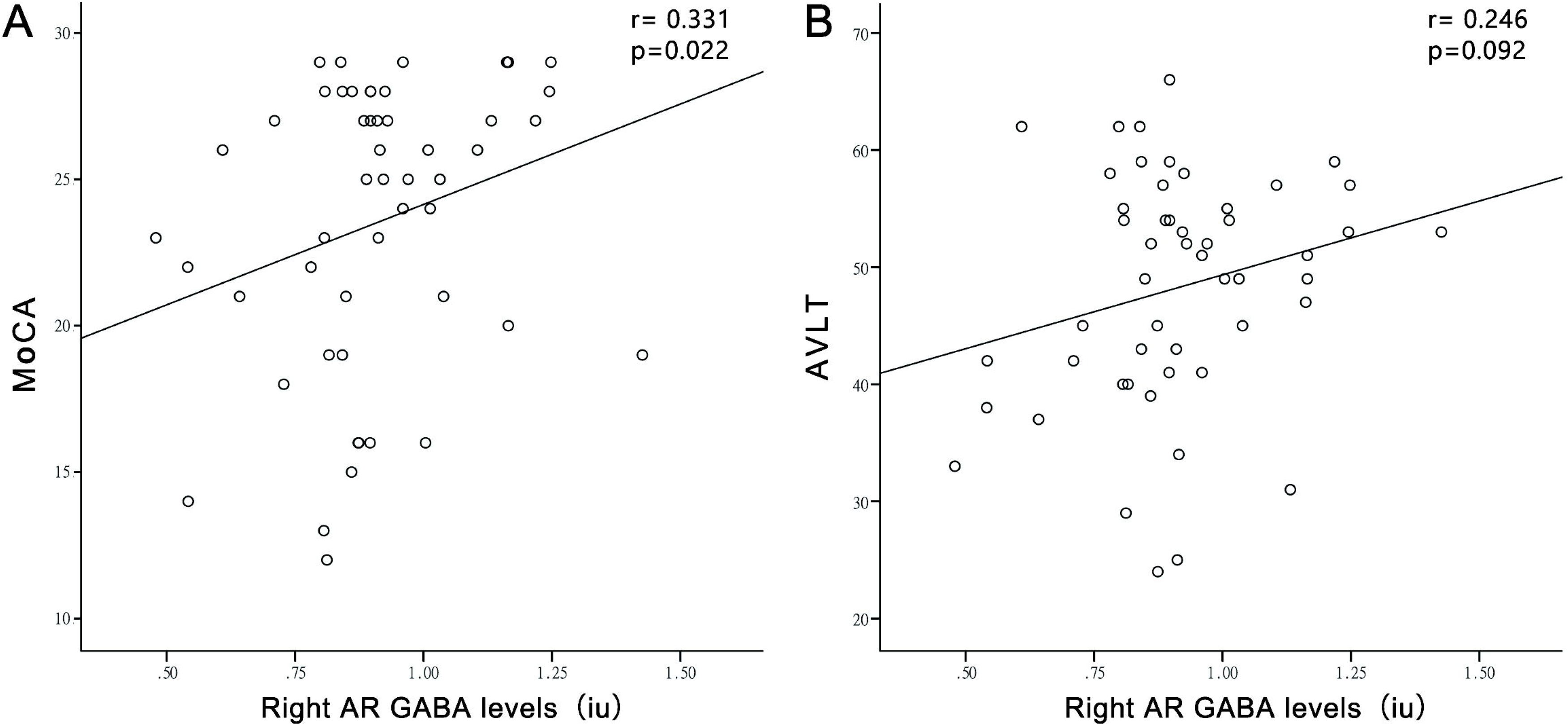
Relationships between GABA/Glu levels and clinical performance in the PC group. GABA levels in the right auditory region (AR) were positively correlated with MoCA (**A**), while the trend of correlation was found between GABA levels in the right AR and AVLT (**B**).

### Feature importance of imaging indexes in predicting clinical features

Based on the random forest model, we evaluated the importance of each feature in terms of IncMSE% and IncNodePurity. To filter out the redundant features, we extracted the features which show the top30 value in IncMSE% and IncNodePurity. Particularly, the prediction ability of MOCA can be significantly increased (paired *t*-test, *p*<0.05) after feature deduction by random forest. Here in Figure 10, we visualize the top 30 features that show the highest contributions to the model. We observed that the imaging measurements that exhibited significant group differences, also show high engagement in predicting MOCA (ranked within top10).

**Figure 10.**
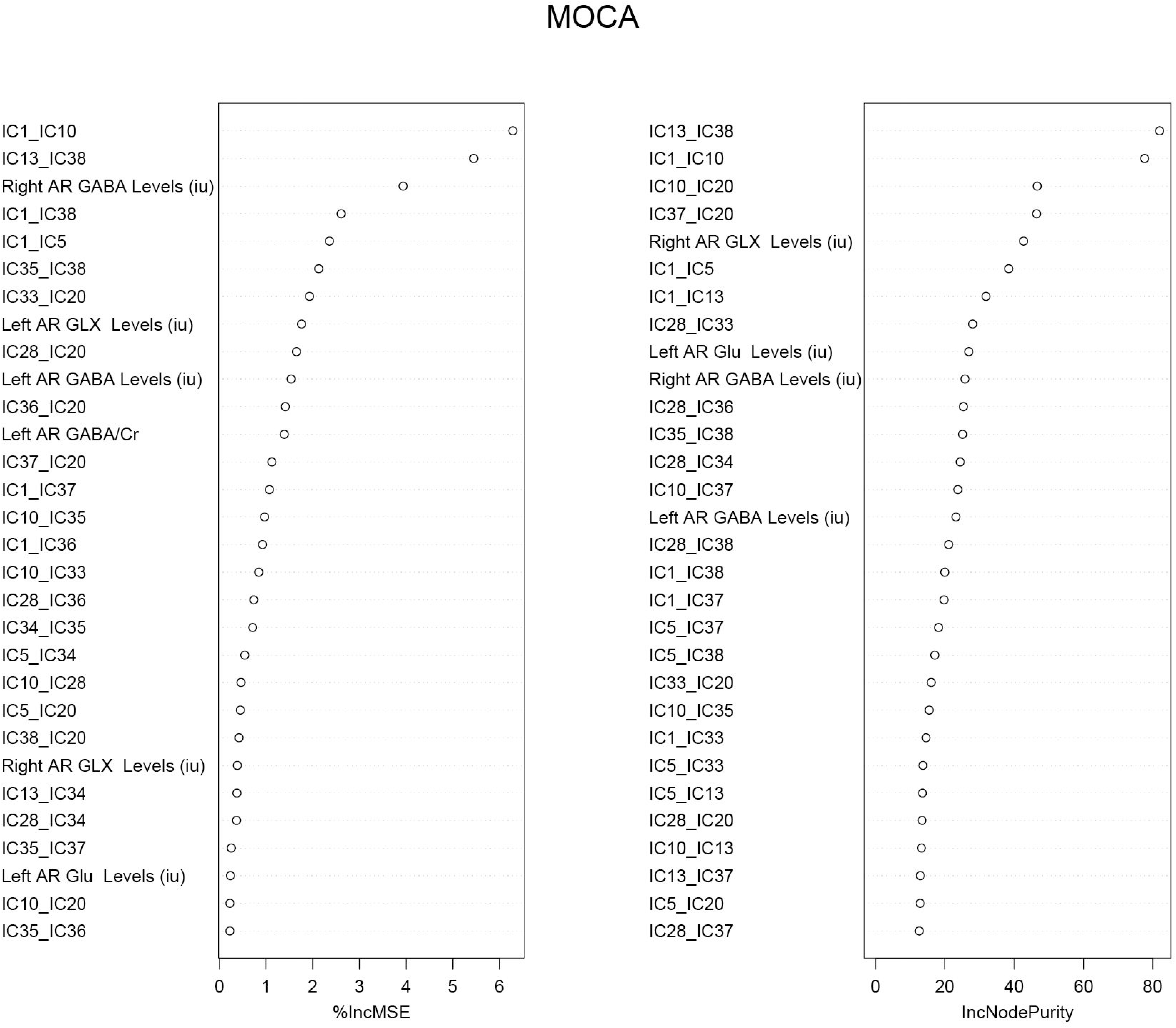
The importance of each feature contributed to MoCA prediction. Top 30 important features are shown in terms of IncMSE and IncNodePurity. IncMSE means the increased percentages of MSE (Mean Squared Error) after rearranging the feature randomly. IncNodePurity means the increased percentages of node purity after rearranging the feature randomly.

## Discussion

As far as we know, this research firstly combined rs-fMRI and edited MRS to investigate changes of intra- and inter-network FC and their relationships with abnormal auditory GABA and Glu levels and CI in PC patients. The main findings include: (1) intra-network FC decline in the DMN and ECN, and inter-network FC decline between AN and DMN and between SN and BGN or VN in PC, as well as increased inter-network FC between aDMN and SN which correlated with hearing loss; (2) Glu levels decline in the bilateral ARs and GABA levels decline in the right AR, and right AR GABA levels correlated with MoCA in PC; (3) the right AR GABA levels correlated with FC strengths between AN and DMN in PC, which associated with executive control; (4) the right AR Glu levels correlated with FC strengths between AN and pDMN in PC, which associated with verbal learning and memory ability. Altogether, these findings suggest neurochemical underpinnings of functional remodeling of cognitive-ear link at the large-scale brain network level, which were closely related to the CI of PC patients.

### Decreased intra-network FC in PC

We found decreased FC of PC patients mainly in bilateral precuneus and PCC within the DMN, which plays a critical role in cognitive processing, shows high activity in rest but is deactivated during stimulus- or task-induced activity ^45, 46^. As an advanced cognitive network, the key nodes of DMN have different function but with high intra network consistency. PCC is the key node of DMN, it has important role in memory and cognition, information processing, and can coordinate auditory stimulation under nonoptimal conditions ^47^. In addition, precuneus mainly participates in the episodic memory and visuospatial processing ^48^. In previous study, we have found reduced cortical volume of the bilateral precuneus and right PCC in PC patients ^14^. Moreover, a recent fMRI study on SNHL patients also found reduced FC related to DMN, mainly including PCC and bilateral precuneus, it was consistent with our study ^49^. In short, PC-related hearing loss may increase listening effort and complete the top-down network control through a series of cognitive compensation, which further affect the consistency within the network, resulting in remodeling of DMN. Interestingly, we also observed a trend of correlations between precuneus FCs and PTA in PC group, which may indicate remodeling of DMN may be dependent on HL to some extent.

Executive control network (ECN) also showed decreased intra-network FC in PC patients in our study. ECN is mainly participated in the executive functions, including work memory, adaptability of cognitive tasks, problem solving and planning ^50^. Decreased FCs were found in the right SFC and dlPFC in rECN, while the left SFC and IPL in lECN. Importantly, SFC FCs obviously correlated the MoCA and SDMT, and the trend of correlation was also found between dlPFC FCs and SDMT. Previous fMRI study has revealed decreased low-frequency fluctuation in the dlPFC, SFC and IPL in PC patients and abnormalities of intra-network FC within right ECN in patients with SNHL, which were well consistent with our results ^15, 49^. The dlPFC is implicated in multisensory integration and top-down regulation processes, and it processes a large amount of cortical information collected through the thalamic reticular nucleus, which was the most important part of auditory pathway ^51^. The SFC, including the supplementary motor area (SMA), mainly participates in speech perception, auditory processing, and auditory representation. The activity enhancement in the SMA had been confirmed in a fMRI study if the voice signal is not clear or in noisy environments ^52^. Moreover, the SMA and IPL also play an important role in accurate control of auditory-motor actions ^53^. Thus, intra-network FC changes in dlPFC, SMA and IPL may induce functional impairment in auditory pathway, auditory processing, and speech function in PC patients, which indicated that long term auditory input impairment may arise cross-modal reorganization. This is especially reflected as speech perception difficulties in noisy environments in patients with PC.

### Altered inter-network FC in PC

Although no changes of intra-network FC were found in AN, a significantly decreased inter-network FC between AN and DMN of PC patients was confirmed in our study. The theory of cognitive-ear link is based on the coupling effects between auditory and cognitive processing ^7^. Our findings suggest decrease auditory cortical stimulation might lead to a bottom-up cascade and result in reduced coupling between AN and DMN, which indicate the functional remodeling of cognitive-ear link and the compromise of dynamic interaction for auditory-cognitive processing because cognitive reserve have been diverted to maintain normal speech perception of PC patients. Moreover, FC strengths between AN-DMN were significantly correlated with executive control, while FC strengths within AN-pDMN were significantly correlated with verbal learning ability and general cognitive function in PC patients, respectively. Thus, this plasticity of cognitive-ear link involving uncoupling for AN-DMN was also associated with negative cognitive outcomes of PC, and these dysconnectivity above may contribute to permanent impaired cognitive function over time.

A series of inter-network FC changes with SN as the core were also found in our study. The SN commonly plays a regulatory role in DMN and central executive network by integrating the internal and external environmental stimulation, including stimulation from the auditory pathway. The dorsal anterior cingulate cortex (dACC) and anterior insular cortex (AI) were two major hubs of SN ^54^. Previous task-based fMRI studies have shown that when additional attention resources are needed to maximize speech comprehension, SN activation in the elderly increases with the acoustic degradation of speech stimuli ^55^. In our study, inter-network FC between SN I (dACC as the core hub) and VN, BGN and aDMN were deceased, although inter-network FC between SN II (AI as the core hub) and aDMN was increased in PC patients. As confirmed, there were wide range of fiber connections between the dACC and limbic system and the frontal and parietal cortex, which plays a key role in cognitive execution associated with learning and memory. dACC can also receive high-processed auditory information from the rostral STG and prefrontal cortex which mainly responsible for balancing the bottom-up and top-down regulation of auditory information ^56, 57^. The imbalances between the SN and other functional networks in this study could potentially disclose the neuropathological mechanism underlying plasticity of cognitive-ear link from another perspective, in which SN plays an important coordinating role. Interestingly, we found the more severe the HL, the closer the FC between SNII and aDMN in our study. Thus, PC-related HL can cause remodeling of higher-order cognitive networks, and contrarily improved hearing condition might alleviate this large-scale functional remodeling. A series of cohort studies had confirmed that to a certain extent the use of auditory amplification technology can alleviate hearing, and then may improve overall cognitive ability and reduce the risk of dementia ^1^.

### Altered auditory GABA and Glu levels in PC

Our study confirmed that GABA levels in the right AR decreased significantly in PC group, while no changes were found in the left AR. Our previous study has demonstrated lower GABA+ levels in the bilateral AR in PC patients using MEGA-PRESS, which partially consistent with current results ^58^. One possible reason for the discrepancy could be that the detected editing GABA signal contains an important contribution of macromolecules co-edited using MEGA-PRESS. A new method for macromolecule suppression which can detect pure GABA was applied in this study ^40^. To our knowledge, this is the first and the largest study using MEGA-PRESS with macromolecular suppression to study the auditory GABA levels in PC. Our findings may reflect the dysfunctional GABAergic neurotransmission in the central auditory system, consisting with accumulated evidences confirmed by animal model of PC, including decreased GABA release, reduced GABA neurons numbers, decreased GABA levels and reduced glutamic acid decarboxylase levels in the auditory cortex ^59–61^. Moreover, GABA levels in the right AR were positively correlated with MoCA in PC patients and have a trend toward correlation with AVLT which evaluates auditory memory and language learning. In line with our results, right auditory GABA levels were correlated with speech-in-noise understanding reflecting central auditory function ^62^, and correlated with neural dedifferentiation, an important predictor of cognitive decline in the elderly ^29^. Therefore, these findings imply that GABAergic inhibition altered in central auditory processing and may play an important role in central mechanisms of CI in PC. A previous MRS study showed that Glu levels in bilateral ARs of PC patients were reduced compared with young controls ^63^. This study lacked age-matched NH controls, hence altered auditory Glu levels may be mainly caused by age rather than PC. We found decreased levels of bilateral auditory Glu in PC patients compared to age-matched controls, indicating reduced glutamatergic excitatory in the central auditory system. Interestingly, a genome-wide association study (GWAS) reported that the variation of glutamate metabotropic receptor 7 (GRM7) may lead to the developing of individual PC by changing the mechanism of glutamate excitotoxicity susceptibility ^64^.

### Relationships between auditory GABA/Glu levels and FC strengths

Neural circuits are largely dependent on the excitation–inhibition balance (EIB), which is closely associated with the activities of glutamatergic and GABAergic neurotransmission ^65^. Specifically, Glu-mediated excitation provides the basis for the generation of BOLD fMRI signals ^66^, and GABA-mediated inhibition modulate the BOLD signal indirectly and maintain synchrony in neural circuits ^44^. In other words, excitatory Glu and inhibitory GABA signals mediate local and long-range neural circuits. In our study, the right auditory GABA levels were positively related to FC strengths between the AN and the DMN in PC patients, which were significantly decreased and associated with TMT-A assessing executive control. Previous *in vitro and in vivo* researches have shown that altered GABA inhibition reduces the range and accuracy of long-range neural network synchronization ^67, 68^. More important, a recent PC animal model study showed that the expression of GABA and metabotropic glutamate receptors related to plasticity in cortex and hippocampus changed widely during the adaptation of cortex to cumulative hearing loss ^69^, which provided new insights into the hypothetical causal basis of CI in PC patients. In short, these results support the hypothesis that partial hearing deprivation might lead to a reduction of GABA mediated inhibition in central auditory processing, which play a key role in a dysconnectivity within AN and DMN, and the dysconnectivity may contribute to CI of PC. Additionally, right auditory Glu levels in PC were positively correlated to FC strengths within the AN and pDMN, which were obviously associated with MoCA and AVLT assessing verbal learning and memory ability. However, between groups showed no significantly differences in the inter-network FC between AN-pDMN in current research, which may be the result of compensation for Glu-mediated excitation.

Several limitations should be considered in our study. First, as cross-sectional research, our study still cannot determine the causal relationship between PC and brain network remodeling, and the effect of PC aggravating over time on brain network remodeling. This will be further elaborated through subsequent longitudinal studies. Second, there were no changes of intra-network FC within AN. We speculated that a potential reason may be that patients with PC were all mild-to-moderate HL in this study. Besides, except for the bottom-up input pathway, there is also a top-down control pathway of the auditory processing, and the cross-modal plasticity in brain structure and function have been confirmed ^70^. Therefore, the compensation of other sensory and cognitive cortices might avoid dysfunctional FC within AN in patients with PC. Exploring the effect of severe HL on the cognitive-ear link in PC needs further study. Third, most GABA is located in two pools within neurons, namely cytoplasm and presynaptic vesicles ^71^. As we know, MRS can only detect the total GABA in the specified local area, but cannot distinguish these individual GABA ^72^. Thus, the field of our research on GABAergic neurotransmission as a target for PC therapy may be limited.

### Conclusion

Our study confirmed the plastic model of cognitive-ear link at the large-scale brain network level, including the dysconnectivity within high-order cognitive networks and between the auditory-cognitive network and overactivation between cognitive networks dependent on hearing loss, which was closely related to the CI in PC patients. Moreover, GABA levels and Glu levels in the central auditory processing were abnormal in PC patients. Importantly, reduction of GABA-mediated inhibition plays a key role in a dysconnectivity between the auditory-cognitive network, which may be neurochemical underpinnings of functional remodeling of cognitive-ear link in PC. Research on GABA neurotransmission regulation may promote the development of new treatment strategies for CI in PC patients.

## Declaration of Competing Interest

None to disclose

## Acknowledgments

This work was supported by the National Natural Science Foundation of China for Young Scholars (No. 81601479), Taishan Scholars Project (No. tsqn201812147), Jinan Science and Technology Development Program of China (Nos. 201907097 and 202019098), and Academic Promotion Programme of Shandong First Medical University (No. 2019QL023). This work was also supported by NIH grant R01 EB016089, R01 EB023963, R21 AG060245, P41 EB015909 and P41 EB031771.

## Notes

### Competing Interest Statement

The authors have declared no competing interest.

