## Supplementary material for "Dysconnectivity between auditory-cognitive network associated with auditory GABA and glutamate levels in presbycusis patients": Material and methods

### Subjects

The exclusion criteria were as follow: (1) ear diseases that affect hearing thresholds and sensorineural hearing losses other than presbycusis (PC); (2) asymmetrical or conductive hearing loss, Meniere's disease, acoustic neuroma, tinnitus, self-reported hyperacusis; (3) history of ototoxic drugs, otologic surgery, head injury or stroke, previous or now noise exposure or hearing aid use; (4) neurological or mental illness (5) Contraindications of MRI.

### Auditory Assessment

Before audiometry, otoscopic examination was applied to remove cerumen and ensure the integrity of tympanic membrane. Then, tympanometry was executed using the Madsen Electronics Zodiac 901 Middle Ear Analyzer to ensure the normal functional status of the middle ear. The pure tone threshold was assessed via a clinical audiometer (Madsen Electronics Midimate 622) coupled with TDH-39P telephonic headphones for each ear separately at frequencies of 0.125, 0.25, 0.5, 1, 2, 4, and 8 kHz. Speech detection was assessed using speech reception threshold (SRT), and Automatic HOPE software was adopted to deliver and evaluate the spondee words in quiet condition. The test was performed according to the SRT guidelines recommended by the American Speech Hearing Association.
